# Differential Control of HIV-1 Replication by IFN-α14 Compared to IFN-α2 Relates to Differences in the Modulation of Host Antiretroviral Restriction Factors

**DOI:** 10.1101/2025.11.01.685890

**Authors:** Saurav S. Rout, Madeline T.H. Stewart, Nathan B. Seidel, Ulf Dittmer, Kathrin Sutter, Kerry J. Lavender

**Affiliations:** Department of Biochemistry, Microbiology and Immunology, College of Medicine, University of Saskatchewan, Saskatoon, SK, S7N 5E5, Canada; Institute of Virology, University Hospital Essen, University of Duisburg-Essen, Essen, Germany

**Keywords:** HIV-1, restriction factor, interferon-alpha, subtypes

## Abstract

Type I IFN, including IFN-α, induces the expression of antiviral restriction factors that can interfere with multiple steps of the HIV-1 replication cycle. Humans have 13 IFN-α genes which encode 12 different IFN-α subtypes. Our previous work in HIV-1 infected humanized mice showed that IFN-α14 treatment more potently controlled HIV-1 than treatment with the clinically approved IFN-α2 subtype. However, the mechanisms behind the more potent control of HIV-1 by IFN-α14 are unknown. The IFN-α14 subtype is known to more potently induce the expression of the restriction factors MX2 and ISG15 and increased APOBEC3G signature mutations *in vivo* compared to IFN-α2. To study the importance of each of these restriction factors in mediating the potent control of HIV-1, we used a CRISPR-Cas9 lentivirus system to create stable knockouts in the MT4C5 cell line that is susceptible to HIV-1 but does not produce measurable amounts of endogenous IFN-α. Knock out of ISG15, but not MX2, eliminated differences in viral suppression after IFN-α14 and IFN-α2 treatment. Similarly, APOBEC3G deletion eliminated differences in viral suppression and the number of infectious particles produced after IFN-α14 and IFN-α2 treatment. Furthermore, APOBEC3G deletion resulted in significantly fewer GG→AG mutations in viral DNA isolated from target cells incubated with supernatant from IFN-α14 treated groups. However, APOBEC3G knock out did not result in significant increases in vDNA compared to the wild type in any experimental group. Overall, elimination of APOBEC3G and ISG15 impaired IFN-α14–mediated suppression of HIV-1, highlighting them as downstream effectors of IFN-α14’s more potent anti-HIV-1 activity.

**IMPORTANCE:** This study uncovers the molecular basis for the more potent antiviral activity of IFN-α14 compared to the clinically used IFN-α2 subtype against HIV-1. Although interferons are known to induce numerous restriction factors, the mechanisms underlying subtype-specific antiviral potency remained unclear. By using CRISPR-Cas9 knockout MT4C5 cell lines, the study identifies ISG15 and APOBEC3G as key effectors mediating IFN-α14’s enhanced suppression of HIV-1 replication. Loss of either ISG15 or APOBEC3G abolished the differential antiviral effect between IFN-α14 and IFN-α2, demonstrating their essential roles in IFN-α14 driven viral restriction. These findings highlight that individual IFN-α subtypes engage distinct downstream pathways and that subtype diversity encodes functional specialization rather than redundancy. Overall, this work advances our understanding of innate immune control of HIV-1 and provides a foundation for developing targeted interferon-based therapies that exploit the unique mechanisms of potent subtypes like IFN- α14.

## INTRODUCTION

Type I interferon (IFN) is a crucial component of the innate immune system, playing an essential role in defending against a wide range of viruses(1). In humans, type I IFN includes 12 IFN-α subtypes along with IFN-β, IFN-κ, IFN-ε, IFN-ο, IFN-τ, and IFN-δ(2). All type I IFN signal through the widely expressed IFN-α receptor (IFNAR)(3), triggering the activation of interferon regulated genes (IRG)(4, 5). Interferon-α plays a role in controlling many viral diseases including HIV-1. Produced primarily by plasmacytoid dendritic cells, IFN-α engages the IFNAR and activates JAK–STAT and MAPK pathways(6), inducing hundreds of IRG that interfere with viral replication at multiple stages, from entry to assembly(7, 8). It is now known that IFN-α subtypes induce IRG, including viral restriction factors, in a differential manner(9, 10, 11, 12). Many antiretroviral restriction factors have been identified that can inhibit HIV-1, including classical and well documented apolipoprotein B mRNA editing catalytic polypeptide-like 3G (APOBEC3G), SAM and HD domain-containing protein 1 (SAMHD1), tetherin, ISG15 and tripartite motif-containing protein 5 (TRIM5α) and those of more recent characterization: MX Dynamin Like GTPase 2 (MX2), serine incorporator (SERINC) 3/5, interferon-induced transmembrane proteins (IFITMs), schlafen 11, and membrane-associated RING-CH (MARCH) 2/8(13, 14, 15, 16, 17, 18, 19, 20, 21).

Restriction factors can impede HIV-1 replication in many ways. For example, APOBEC3G is incorporated into HIV-1 virions from a producer cell and exerts its antiviral activity in newly infected cells. The antiviral activity of APOBEC3G includes inhibition of reverse transcription, impairment of proviral integration and deamination of the viral RNA resulting in G to A hypermutation of the nascent viral DNA(22, 23, 24) that can result in defective provirus. The MX2 protein inhibits HIV-1 by interfering with the nuclear import of the viral pre-integration complex, which is a crucial step after viral DNA synthesis but before its integration into the host cell’s genome(25, 26). Additionally, MX2 inhibits viral capsid uncoating, thereby blocking HIV-1 infection(27). ISG15, a ubiquitin-like protein, is the most highly expressed IRG(28). It inhibits both the assembly and release of HIV-1 virions from cells(29, 30).

It has been well documented that IFN-α subtypes differentially control HIV-1 replication. IFN- α14 is the more potent subtype both *in vitro* and *in vivo* when compared to clinically used IFN- α2(12, 31). However, whether the more potent control of HIV-1 by IFN-α14 is due to differential modulation of restriction factors is unknown. Lavender et al. and Harper et al.(31, 32) showed that transcription of *MX2* was significantly increased by IFN-α14 but not IFN-α2. In contrast, *APOBEC3G* was not transcriptionally upregulated but there was a significant increase in APOBEC3G-mediated signature mutations in the proviral HIV-1 DNA (vDNA) from IFN-α14 compared to IFN-α2 treated mice. In 2021, Tauzin et al.(12) showed that IFN-α14 also induced greater *ISG15* expression in MT4C5 cells. Since different IFN-α subtypes appear to modulate different restriction factors and control HIV-1 in a differential manner, we wanted to investigate whether the MX2, ISG15 and APOBEC3G restriction factors are important in IFN-α14’s more potent control of HIV-1. We used a CRISPR-Cas9 system to knock out each restriction factor in MT4C5 cells to study their importance in the more potent control of HIV-1 by IFN-α14 compared to IFN-α2.

## RESULTS

### IFN-α14 more potently reduced HIV-1 replication than IFN-α2 when used at a biologically relevant activity

The relationship between protein concentrations and the functional activity of cytokines such as IFN-α is not strictly linear, as functional activity is influenced not only by expression levels but also by factors such as folding efficiency when using recombinant proteins and ligand-receptor interactions. Different IFN-α subtypes have been shown to bind with different affinities and kinetics to the interferon alpha receptor (IFNAR)(33, 34, 35, 36, 37). For example, the IFN-α14 subtype binds more tightly to IFNAR compared to IFN-α2(38) and this difference in affinity could potentially influence the intensity of the downstream signaling cascade(39) leading to differential expression of IRGs. Also, doses of biologics like interferons should be expressed in activity units rather than mass units to account for the presence of inactive protein fractions in recombinant formulations(40). Since the IFN-α2 and -α14 subtypes in our study were produced recombinantly and have different affinities for IFNAR, we first needed to normalize their concentrations based on signaling activity in order to better understand their potentially different antiviral effects(31, 41). To normalize the activity between IFN-α2 and IFN- α14 we used a reporter cell line containing an interferon-stimulated response element (ISRE) promoter/enhancer element driving a luciferase reporter gene. Using the reporter cell line, we titrated each IFN-α subtype over a range of protein concentrations and normalized the activities of each subtype based on their EC50 values to that of an IFN-α standard with a known activity prior to use in experiments (Fig. 1A).

**Figure 1.**
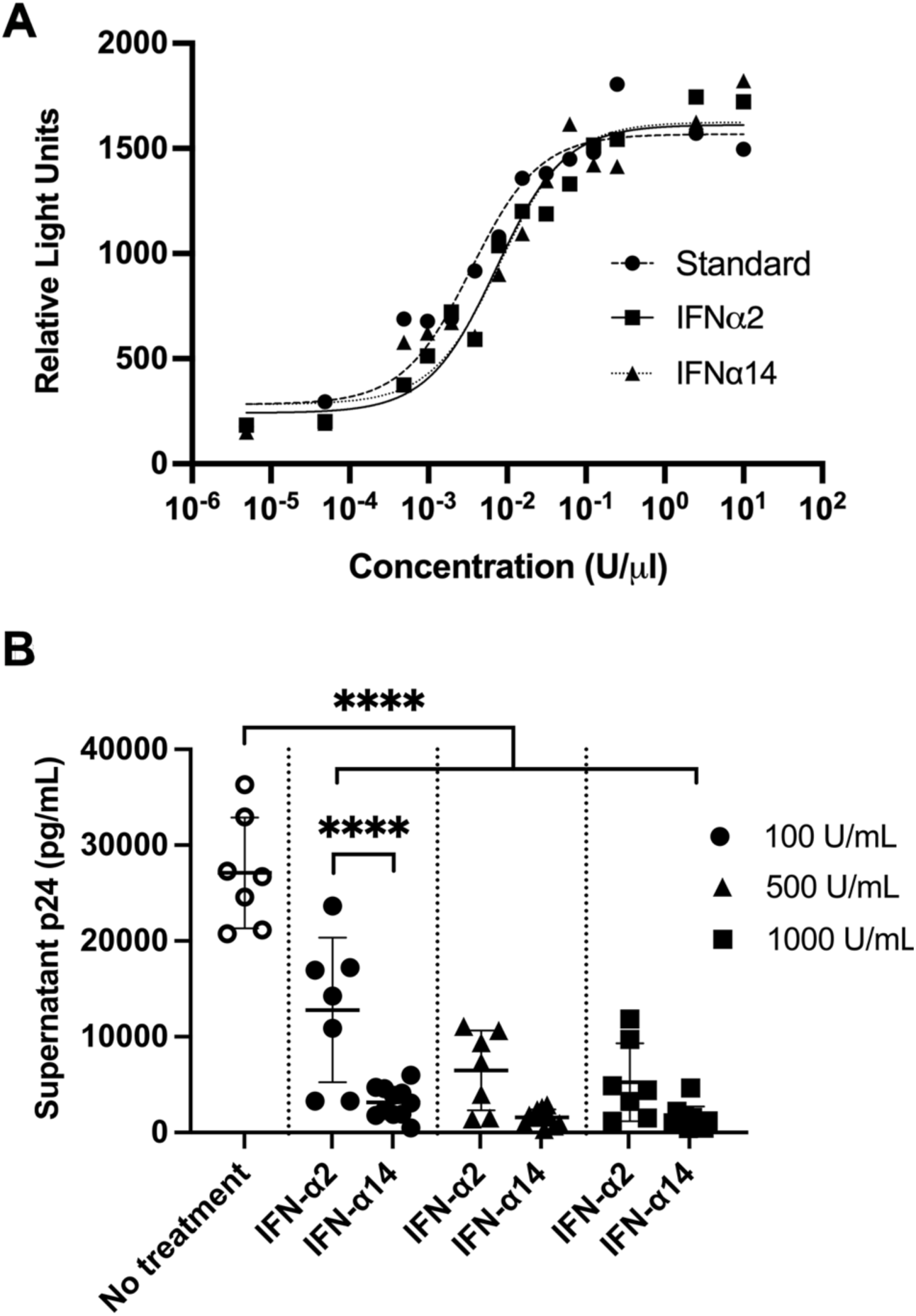
IFN-α14 more potently suppressed HIV-1 replication than IFN-α2 at similar activity levels. (A) The activities of IFN-α2 and IFN- α14 were determined by titration of each subtype on an IFN-α responsive reporter cell line in comparison to an IFN-α stock of known activity. (B) HIV-1 p24 ELISA of supernatants collected from MT4C5 cells treated with 0, 100, 500 or 1000U/ml of IFN-α2 or IFN-α14 for 6 days starting 2 hours post infection. One-way ANOVA with Tukey’s multiple comparison test. **** p<0.0001

To evaluate the importance of antiretroviral factors in mediating more potent control of HIV- 1 by IFN-α14 compared to IFN-α2 we utilized the MT4C5 cell line. The MT4C5 cell line is a derivative of MT4 cells that additionally expresses CCR5 and can be infected by CXCR4-tropic (X4) or CCR5-tropic (R5) HIV-1 viruses. Importantly, following HIV-1 infection, MT4C5 cells do not produce measurable amounts of endogenous IFN-α in the cell culture supernatant(42) that may otherwise confound findings. We first established our system in MT4C5 cells to test the contributions of each restriction factor to the more potent control of HIV-1 by IFN-α14. Wild-type (WT) MT4C5 cells were infected with HIV-1JR-CSF at 0.1 MOI for 2 hours and then treated with a biologically relevant activity (100U/ml)(43) or two supraphysiological doses (500U/ml or 1000U/ml)(44) of IFN-α2 or IFN-α14. Three days post infection, fresh medium with 100U/ml, 500U/ml or 1000U/ml of IFN-α2 or IFN-α14 was replenished. Six days post infection, supernatant was collected and a p24 ELISA performed to quantify HIV-1 replication through the production and release of HIV-1 virions into the supernatant. Both IFN-α subtypes significantly reduced p24 in supernatants (Fig. 1B). However, supernatants from cells cultured in 100U/ml of IFN-α2, contained significantly higher amounts of viral p24 compared to cells cultured in 100U/ml of IFN-α14 (Fig. 1B). In cultures containing 500U/ml and 1000U/ml of IFN-α, there were no significant differences observed between subtypes, although a trend toward lower levels of virus production could be detected in the cultures containing IFN-α14 (Fig. 1B). The results indicated that greater suppression of HIV-1 replication in MT4C5 cells by IFN-α14 compared to IFN-α2 occurred at the biologically relevant activity(43) of 100U/mL, but less so at supraphysiological concentrations.

### Potent suppression of HIV-1 replication by IFN-α14 occurs intracellularly

We first investigated if lower levels of virus in supernatants from IFN-α14 treated cells might have resulted from an accumulation of virions within the cell with inhibition of HIV-1 release into the supernatant. We performed a p24 immunoblot using cell lysates from untreated HIV-1 infected MT4C5 cells and cells treated for 6 days with 100U/ml of either IFN-α2 or -α14. The immunoblot indicated that lysates from cells treated with IFN-α14 had no detectable cell-associated p24 whereas p24 was clearly visible in lysates from cells treated with IFN-α2 (Fig. 2A). Furthermore, quantification of the p24 bands in relation to the GAPDH loading control on the western blot confirmed there was some reduction in cellular p24 in the cells treated with 100U/ml of IFN-α2 compared to untreated cells as expected based on the p24 ELISA results in Fig. 1B, although not nearly to the degree as in IFN-α14 treated cells (Fig. 2B). The results indicated that the more potent suppression of HIV-1 replication by IFN-α14 than IFN-α2 at the biologically relevant concentration of 100U/mL(43) did not primarily originate due to accumulation of cell-associated virions that were not being released into the supernatant.

**Figure 2.**
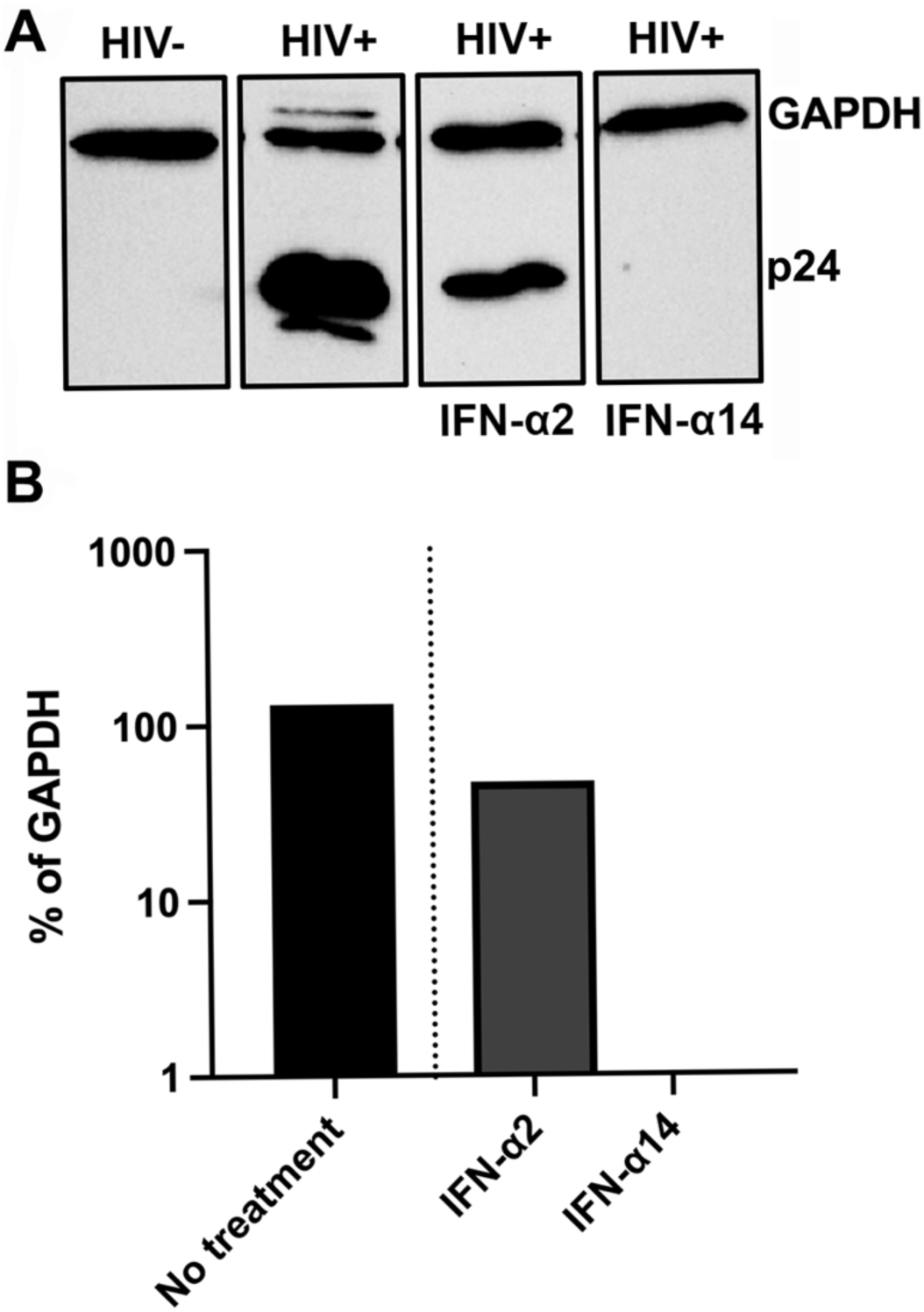
IFN-α14 more potently reduced cellular p24 levels in MT4C5 cells than IFN-α2. (A) HIV-1 p24 immunoblot of MT4C5 cell lysates previously treated for 6 days with IFN-α2 or IFN-α14. GAPDH loading control. (B) Quantification of the p24 immunoblot normalizing the intensity of the p24 bands to GAPDH.

### IFN-α14 treatment more potently reduced cellular HIV-1 viral DNA (vDNA) and viral infectivity compared to IFN-α2

Having established that IFN-α14 potently suppressed levels of HIV-1 p24 within the cell we next evaluated whether the reduction in p24 was due to reductions in viral HIV-1 DNA (vDNA) within cells and/or the amount of infectious virus being produced over the 6-day infection. First, we evaluated levels of vDNA in HIV-1 infected MT4C5 cells that had been left untreated or treated with 100U/ml of either IFN-α2 or IFN-α14 for 6 days. To quantify the levels of vDNA we isolated genomic DNA from the cells at day 6 and performed qPCR targeting the HIV-1 POL region. We found that cells treated with 100U/ml of IFN-α14 contained relatively lower levels of total vDNA compared to untreated and IFN-α2 treated cells (Fig. 3A). Cellular input levels were normalized to copies of *GAPDH.* Reductions in cellular p24 and vDNA over 6 days of infection could also relate to antiretroviral restriction factor-mediated production and release of defective virions. Therefore, we assessed if IFN-α14 treatment of MT4C5 cells resulted in the production of fewer infectious HIV-1 virions. Supernatants from cells treated with 100U/ml of IFN-α2 or IFN-α14 were isolated and tested for levels of infectivity per pg of p24 using TZM-bl reporter cells, which contain a Tat-sensitive LTR promotor driving the β- galactosidase gene. Individually infected cells can thus be quantified through the presence of β- galactosidase activity. The infectivity of virus-containing supernatants isolated from cells treated with 100U/ml IFN-α14 was significantly lower than the infectivity of HIV-1 in supernatants after IFN-α2 treatment (Fig. 3B). The data show that IFN-α14 more potently reduced both HIV-1 vDNA and viral infectivity than IFN-α2 in MT4C5 cells.

**Figure 3.**
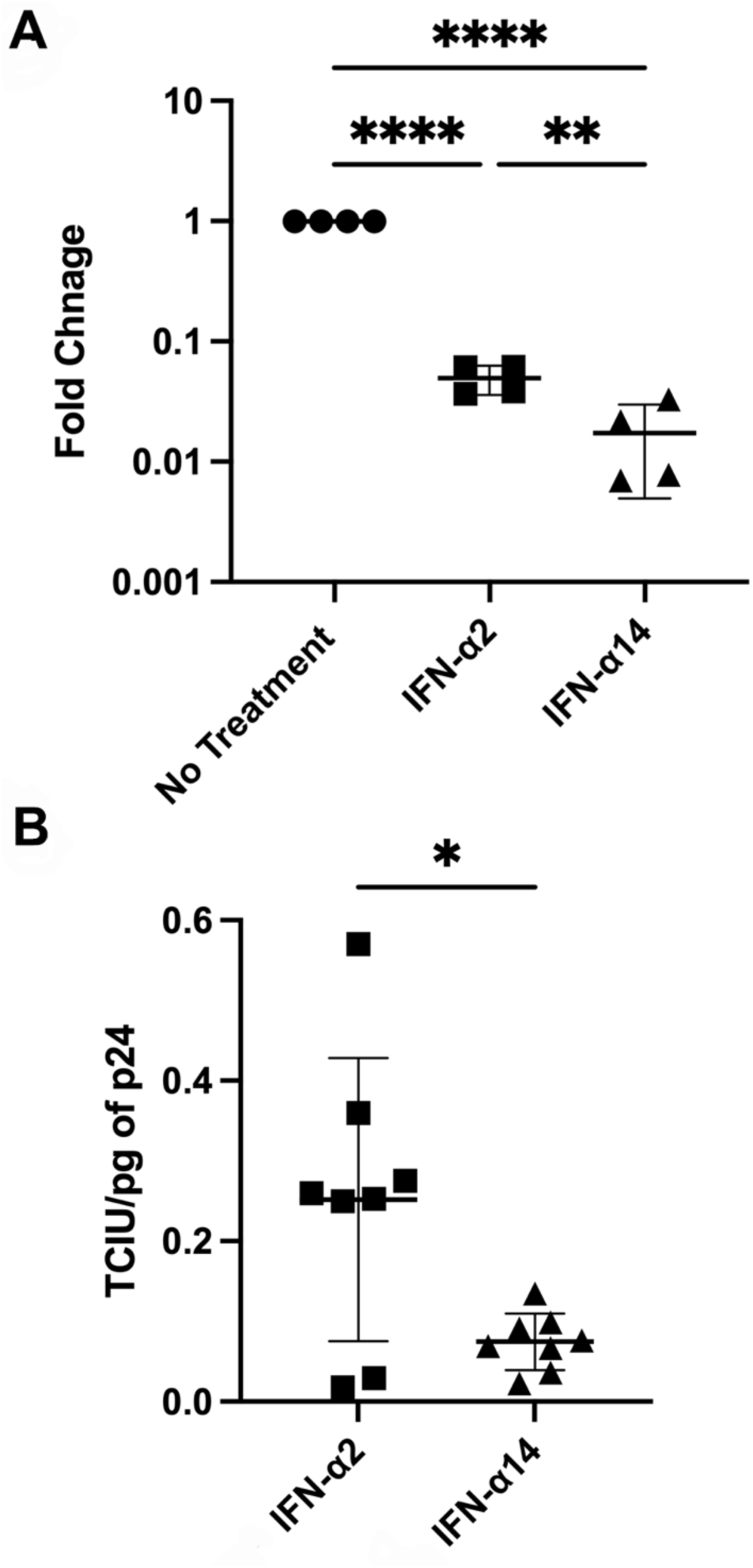
IFN-α14 more potently reduced HIV-1 viral DNA (vDNA) and HIV-1 infectivity in MT4C5 cells than IFN-α2. (A) Fold change in HIV-1 vDNA levels in MT4C5 cells differentially treated with IFN-α upon amplification of the POL region from isolated genomic DNA. Cellular input was normalized to *GAPDH*. One-way ANOVA with Tukey’s multiple comparison test ** p<0.01; **** p<0.0001. (B) Tissue culture infectious units (TCIU) per pg of input HIV-1 p24 was determined by TZM-bl infectivity assay using supernatants from MT4C5 cells treated with 100U/mL of IFN-α2 or IFN-α14. Student’s unpaired t-test * p<0.05.

### CRISPR-Cas9-mediated knock out of MX2, APOBEC3G and ISG15 in MT4C5 cells

At this point in our investigations the antiretroviral restriction factors MX2, APOBEC3G and ISG15 previously shown to be either more highly expressed or activated by IFN-α14, remained as candidates factors mediating IFN-α14’s more potent control of HIV-1 infection(12, 31, 32). To investigate the role of each factor further, we used the CRISPR-Cas9 method to knock out each restriction factor in MT4C5 cells. Briefly, two to three guide oligos were designed for each gene (Table 1) and individually cloned into a lentiCRISPRv2 backbone. Lentiviral vectors carrying the guide sequences were then made by transfecting lentiCRISPRv2, psPAX and pMD2.G plasmids into HEK293FT cells. The MT4C5 cells were then transduced with the each of the resultant lentiviruses for 48h before plating them into selection medium. Successful MX2 and ISG15 gene knock out (KO) was determined using immunoblots and APOBEC3G KO confirmed by PCR due to a specific antibody being unavailable. A concentration of 1000U/ml of IFN-α2 was needed to induce detectable levels of MX2 in WT MT4C5 cells whereas MX2 could be detected in cells treated with either 100 or 1000U/ml of IFN-α14, albeit less so after stimulation with 100U/ml of the cytokine. However, MX2 expression was not detected after stimulation with either IFN-α in the MX2 KO cells (Fig. 4A). To detect ISG15 protein in MT4C5 cells 1000U/ml of both IFN-α subtypes was used. However, no ISG15 was detected in KO cells after IFN-α stimulation (Fig. 4B). The APOBEC3 family is highly homologous at the protein level and thus there are no commercially available antibodies that could specifically detect the APOBEC3G KO in MT4C5 cells. Thus, we used APOBEC3G-specific sequences in genomic DNA to detect the APOBEC3G KO. Amplification of genomic DNA using APOBEC3G sequence-specific primers showed amplification of a shorter band in APOBEC3G KO cells compared to genomic DNA from WT MT4C5 cells (Fig. 4C). In summary, we were able to successfully knock out MX2, ISG15 and APOBEC3G in MT4C5 cells in order to further evaluate the contribution of each antiretroviral restriction factor to the more potent control of HIV-1 infection by IFN-α14 compared to IFN-α2.

**Figure 4.**
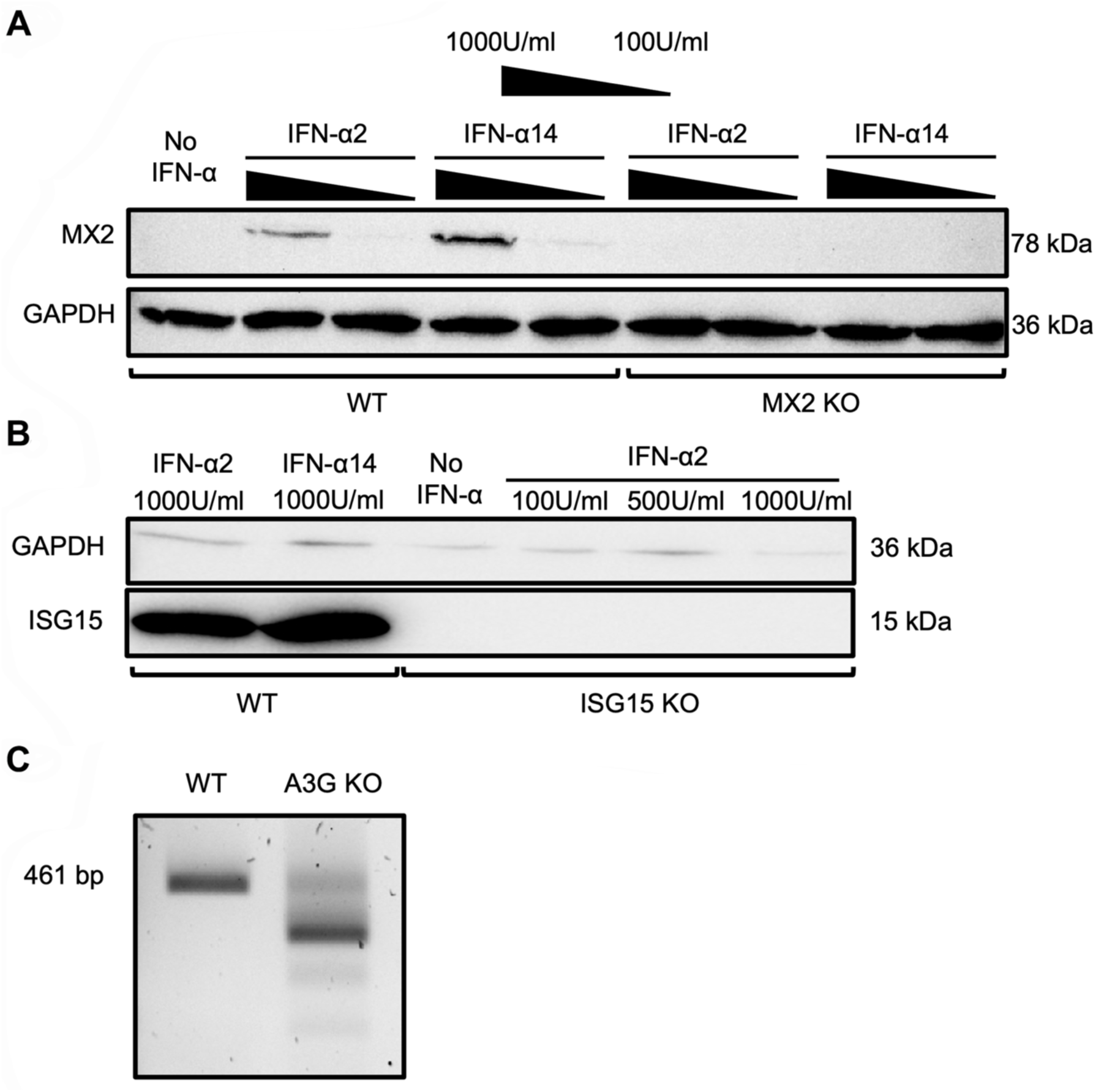
The MX2, APOBEC3G and ISG15 antiretroviral restriction factors were successfully knocked out of MT4C5 cells using CRISPR-Cas9. MT4C5 cell lysates of IFN-α treated wild-type (WT), MX2 knockout (KO) and ISG15 KO cells were run on a 14% SDS PAGE gel. The proteins were then transferred to a nitrocellulose membrane and immunoblotted for (A) MX2 and (B) ISG15. GAPDH was used as a loading control. (C) PCR amplification product of a 461bp region of APOPEC3G (A3G) expected to be disrupted in A3G KO cells compared to WT using APOBEC3G sequence specific primers.

**Table 1.**
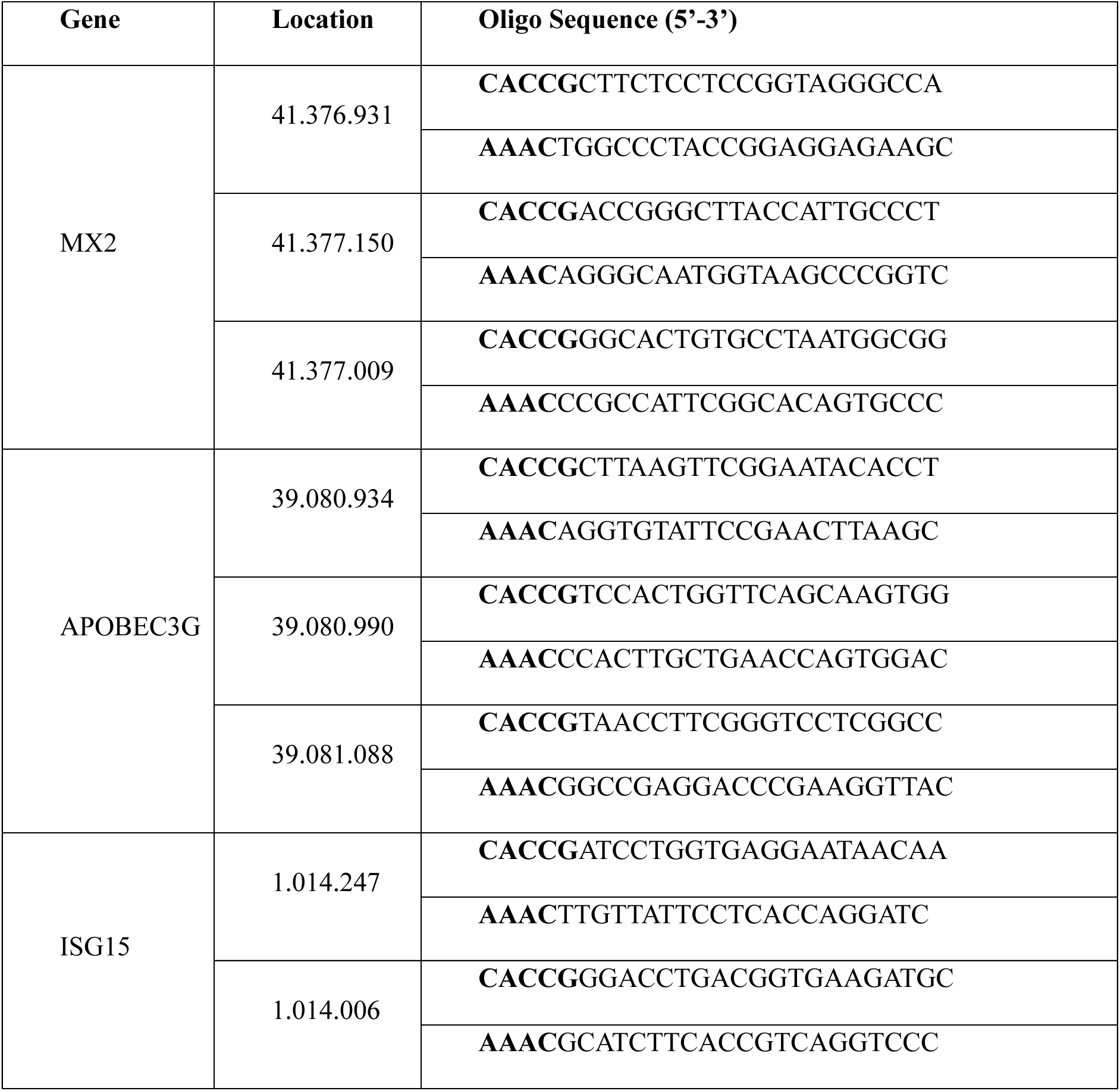
Guide DNA oligo sequences.

### Deletion of APOBEC3G and ISG15 but not MX2 eliminated IFN-α14’s more potent control of HIV-1 compared to IFN-α2

We next used our MX2, APOBEC3G and ISG15 KO cells to evaluate their importance in IFN-α14’s more potent control of HIV-1 compared to IFN-α2. To evaluate whether IFN-α14’s potency was dependent on APOBEC3G, ISG15 or MX2 each of the KO cells as well as WT MT4C5 cells were treated with 100U/mL of either IFN-α2 or IFN-α14 as performed earlier. As expected, IFN-α14 reduced levels of HIV-1 p24 in the cell supernatants compared to IFN-α2 in the WT MT4C5 cells. Unexpectedly, IFN-α14 still reduced p24 levels in supernatants significantly better than IFN-α2 in the MX2 KO cells (Fig. 5). However, supernatants collected from APOBEC3G KO and ISG15 KO cells no longer contained different amounts of p24 regardless of which IFN-α treatment they received (Fig. 5). Therefore, the data suggested that the restriction factors APOBEC3G and ISG15 have a role in the more potent control of HIV-1 by IFN- α14 compared to IFN-α2, whereas MX2 does not.

**Figure 5.**
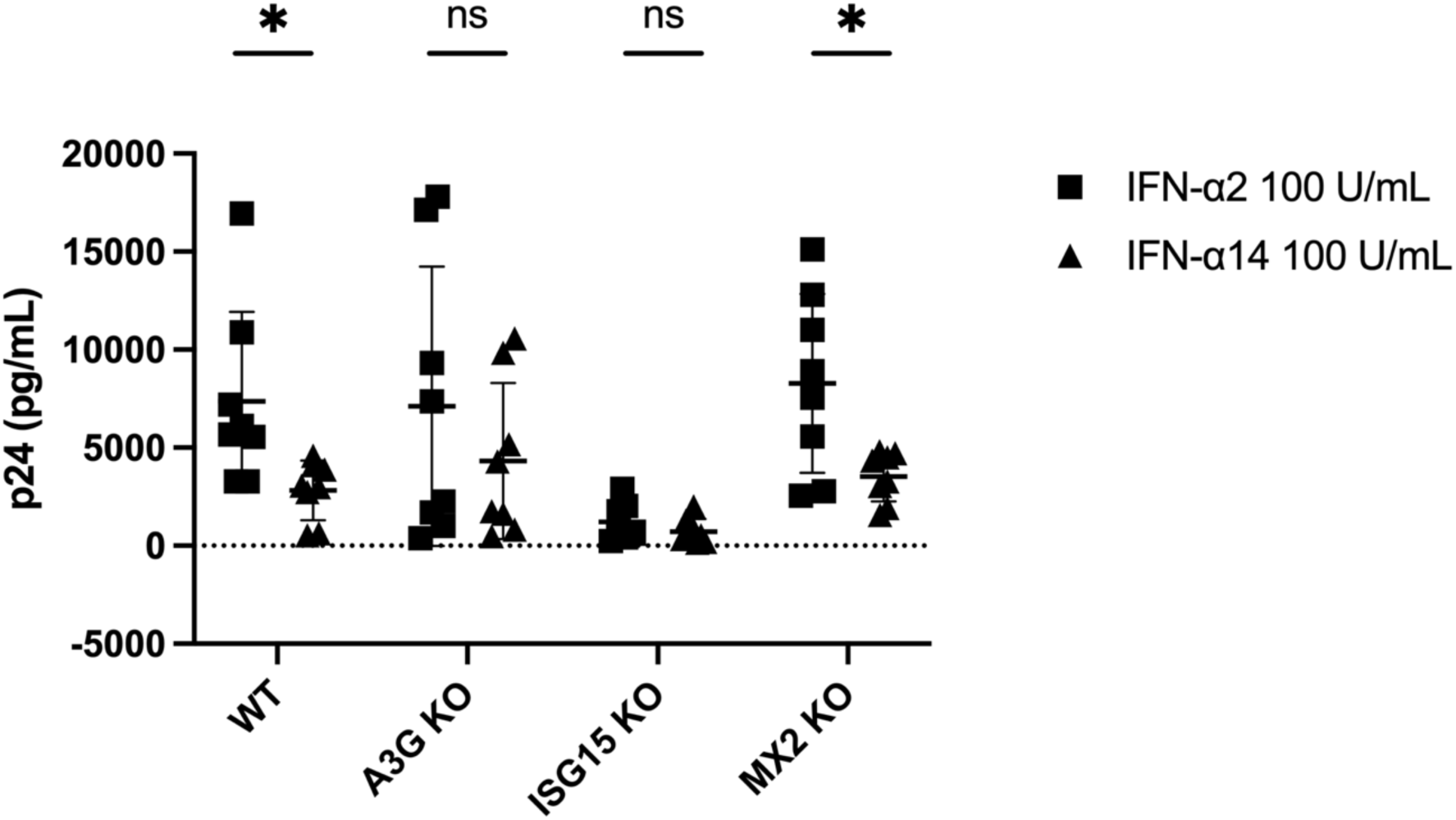
Knock out of APOBEC3G (A3G KO) and ISG15 in MT4C5 cells eliminated differential control of HIV-1 by IFN-α subtypes. HIV-1 p24 ELISA of supernatants collected from HIV-1 infected MT4C5 (WT), A3G KO, ISG15 KO and MX2 KO cells after 6 days of treatment with either 100U/ml of IFN-α2 or IFN-α14. Student’s unpaired t-test * p<0.05.

### IFN-α14 more potently augmented APOBEC3G cytidine deaminase activity and reductions in viral infectivity than IFN-α2 but did not affect vDNA levels

Since HIV-1 vDNA levels and viral infectivity were significantly reduced by IFN-α14 but not IFN-α2 treatment in MT4C5 WT cells, we tested whether these effects were being mediated through the restriction factor APOBEC3G. First, we assessed levels of vDNA in HIV-1 infected MT4C5 WT or APOBEC3G KO cells treated for 6 days with either 100U/ml IFN-α14 or IFN-α2 by amplifying the HIV-1 POL region from isolated genomic DNA as already described. Fold change was calculated from levels of vDNA in untreated MT4C5 WT cells. As we had observed previously, IFN-α14 treatment resulted in significantly lower amounts of vDNA in cells compared to IFN-α2 in MT4C5 WT cells. However, knock out of APOBEC3G did not alleviate the effect (Fig. 6A). In contrast, when we collected day 6 supernatants from MT4C5 WT and APOBEC3G KO cells and evaluated the infectivity of HIV-1 per pg of p24 we observed that APOBEC3G KO restored the infectivity of HIV-1 virions produced from the IFN-α14 treated cells to comparable levels as IFN-α2 treated cells (Fig. 6B). Since the cytidine deaminase activity of APOBEC3G can contribute to reductions in viral infectivity through deleterious mutagenesis of the vDNA we also assessed the frequency of GG → AG signature mutations. We evaluated day 6 supernatants from MT4C5 WT and APOBEC3G KO cells to induce signature mutations in target cells. Interestingly, the frequency of GG → AG signature mutations in the target cells incubated with supernatants from MT4C5 WT cells treated with IFN-α2 was not altered in the absence of APOBEC3G. Conversely, when treated with IFN-α14, knock out of APOBEC3G resulted in a significant reduction in the frequency of signature mutations in target cells (Fig. 6C). Overall, IFN-α14 more potently controlled HIV-1 than IFN-α2 through augmentation of APOBEC3G cytidine deaminase activity leading to the production of more non-infectious HIV-1 virions.

**Figure 6.**
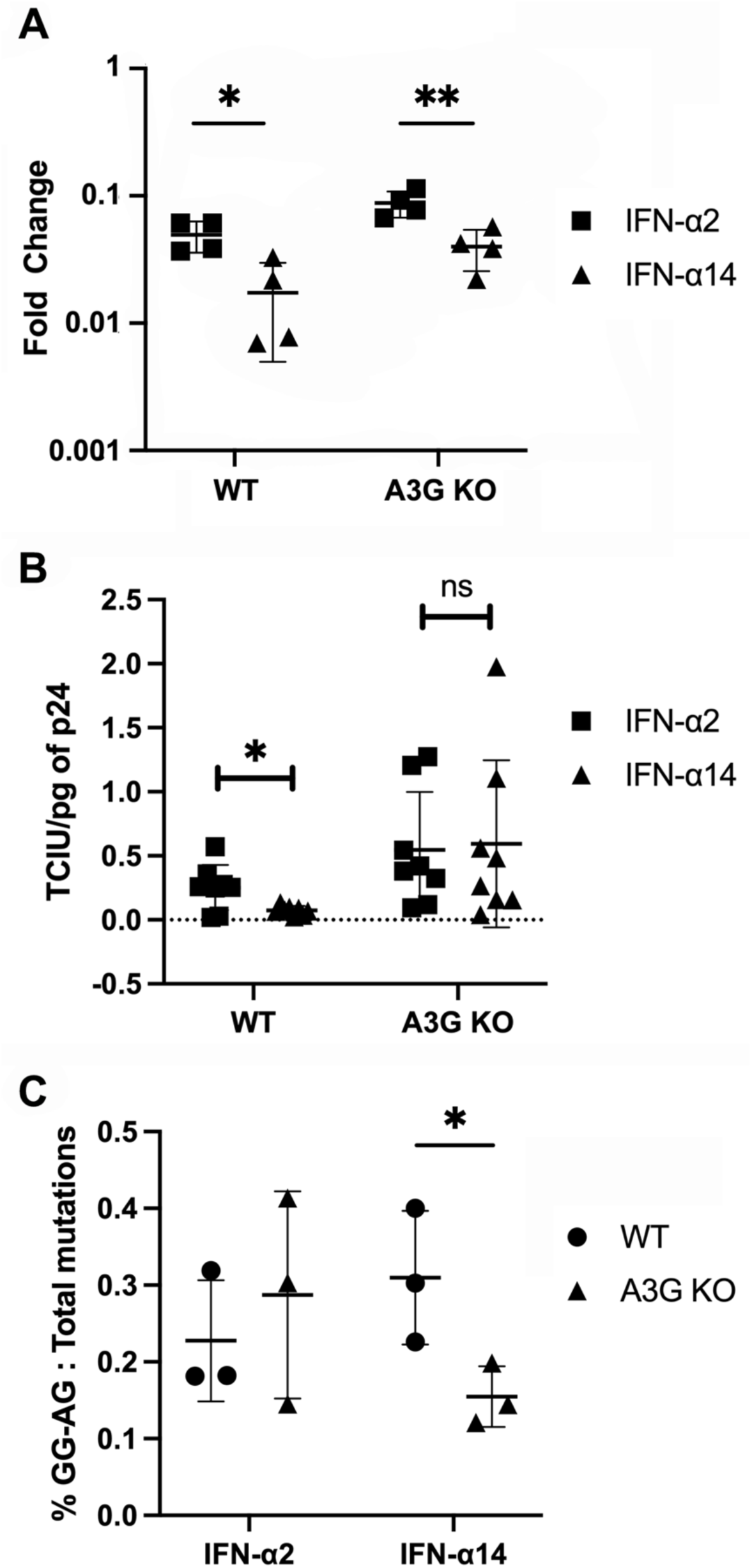
Deletion of APOBEC3G did not alter vDNA in cells but restored infectious particles and frequency of GG → AG signature mutations. (A) Fold change in HIV-1 vDNA levels in MT4C5 WT and A3G KO cells differentially treated with IFN-α upon amplification of the POL region from isolated genomic DNA. Cellular input was normalized to *GAPDH*. (B) TCIU per pg of input HIV-1 p24 was determined by TZM-bl infectivity assay using supernatants from MT4C5 WT and A3G KO cells treated with 100U/mL of IFN-α2 or IFN-α14. (C) Amplified POL region from vDNA of target cells were amplified and sequenced and GG→AG mutations were plotted against total mutations. Student’s unpaired t-test * p<0.05; ** p<0.01.

## DISCUSSION

Greater expression of restriction factors such as *ISG15*, and *MX2* and increased activity of APOBEC3G in IFN-α14 treated humanized mice(12, 31) suggested that IFN-α14’s potency may be, at least in part, due to the contribution of specific restriction factors. The data presented in the current study indicated that the potent HIV-1 suppression mediated by IFN-α14 is driven, at least in part, cell intrinsically by ISG15 and APOBEC3G. We observed that the MT4C5 cells harboring genomic deletion of *ISG15* and *APOBEC3G* showed no difference in viral p24 production when treated with 100U/mL of IFN-α2 or IFN-α14 suggesting that ISG15 and APOBEC3G are responsible for the more potent control of HIV-1 by IFN-α14. It further demonstrated that IFN- α14 treatment not only reduced viral load but also resulted in the production of fewer infectious particles. However, when we increase the dosage of both IFN-αs, the difference ceased to exist. A study done by Schlaepfer and Fahrny et al. showed that at higher doses of IFN-α subtypes, reporter gene induction in response to different IFN-α subtypes waned(45). One of the reasons could be that at higher doses of IFN-α, the IFNAR1/2 receptors become saturated, meaning nearly all receptor complexes are occupied regardless of subtype affinity. When that happens, downstream signaling (e.g., STAT phosphorylation, ISG induction) and antiviral effects may reach a maximal plateau, minimizing observable subtype-specific differences(46). Therefore, we used a biologically relevant dose of 100U/ml(43) where differences in IFN-α responses between subtypes become detectable.

IFN-α14 suppressed virus replication in HIV-1 infected MT4C5 cells. Although p24 was detectable in the supernatant, a complete loss of p24 was observed in the cell lysates of the IFN- α14–treated cells by day 6. This indicated that HIV-1 replication initially occurred following infection, leading to viral production in the supernatant, but IFN-α14 treatment subsequently slowed within cells over the 6 days. This was confirmed by amplifying the POL region of HIV-1 from genomic DNA isolated from cells. However, given that ELISA and PCR are more sensitive than western blot, the absence of p24 detection by western blot does not rule out the presence of cellular virions.

Restriction factors like MX2, APOBEC3G and ISG15 hinder HIV-1 replication at different stages of HIV-1’s lifecycle. MX2 and APOBEC3G act during earlier stages whereas ISG15 acts during the later stages of HIV-1 replication. Since we saw a decrease in viral infectivity in the IFN-α14 treated group compared to IFN-α2 we anticipated that the decrease in infectivity could be a result of increased APOBEC3G activity, therefore we investigated APOBEC3G more thoroughly. ISG15 on the other hand, is known to hinder HIV-1 by preventing ubiquitination of Gag and Tsg101 which is important for viral budding and release(47). ISG15 also modifies a broad range of cellular proteins, many of which are key components of the antiviral innate immune signaling pathway(48). In future, experiments could be performed to determine if ISG15 is restricting HIV- 1 more potently by preventing ubiquitination of Gag and Tsg101 or by modifying other cellular proteins after IFN-α14-treatment. A previous study(31), along with our own findings, concluded that APOBEC3G activity is enhanced during IFN-α14 treatment. However, the mechanism underlying the increased frequency of GG → AG hypermutations in the absence of elevated APOBEC3G expression levels remains unclear. Others have shown that treatment with IFN-α increases *APOBEC3G* mRNA levels 36 folds in a different cell line(49). However, in MT4C5 cell line there was no elevation in *APOBEC3G* mRNA levels(12). Deletion of APOBEC3G did result in comparable levels of HIV-1 cell supernatants between IFN-α treatment groups suggesting that APOBEC3G has a role in IFN-α14’s increased anti-HIV-1 potency. Apart from APOBEC3G, deletion of ISG15 eliminated subtype specific differences in viral load in MT4C5 cells. However, deletion of MX2 did not eliminate the subtype specific differences in HIV-1 control in cells.

A study showed that inactivation of multiple ISGs induced by IFN-β revealed a polygenic and synergistic mechanism underlying HIV-1 control(**50**). Notably, deletion of MX2 did not reduce IFN-α14’s capacity to suppress HIV-1 replication, suggesting that other IFN-α14 induced restriction factors can maintain antiviral activity. In polygenic systems, the effect of one protein can be influenced by the abundance or activity of others, depending on their mode of interaction. ISG15 functions late in the viral life cycle, blocking assembly and release(29, 30), whereas APOBEC3G and MX2 act at earlier stages. When early acting factors such as APOBEC3G or MX2 substantially limit the production of competent virions, ISG15 has fewer targets, making its impact appear smaller; however, when early restrictions are weak, ISG15’s role in blocking release becomes more critical. Therefore, deletion of ISG15 may lead to a greater increase in viral replication compared to deletion of MX2 under IFN-α14 treatment, as the antiviral effect of MX2 can be partially compensated by ISG15, whereas loss of ISG15 is less likely to be offset by other restriction factors. It is plausible that the expression level of a compensatory restriction factor influences HIV-1 replication. However, in our experiments, we did not measure the expression of other restriction factors when a single factor was knocked out during HIV-1 infection. Future studies involving the simultaneous knock out of ISG15, APOBEC3G, and MX2 could help reveal synergistic interactions among restriction factors regulated by IFN-α14. Since the MT4C5 cell line does not produce endogenous IFN-α, it served as a useful model to examine the exogenous effects of IFN-α subtypes. However, not all IRG responses could be evaluated, as MT4C5 cells may respond differently from primary CD4+ T cells. Therefore, future validation using primary CD4+ T cells would be useful to assess the physiological relevance of IFN-α subtype responses.

Here we demonstrated that IFN-α14, in contrast to IFN-α2, exerted a stronger effect on HIV- 1 replication and restriction factors like ISG15 and APOBEC3G play significant roles in the differential effect between IFN-α2 and IFN-α14. However, the contribution of other restriction factors like MX2 and BST2 cannot be ignored in the context of HIV-1 replication. Although in our experimental setup these restriction factors (MX2 and BST2) may seem redundant, but their contribution to control HIV-1 cannot be disregarded. In future, short-term experiments for 2 or 3 days with a fully established infection and IFN-α treatment could be done to evaluate the effect of MX2 and BST2 on IFN-α14 potency.

## MATERIALS AND METHODS

### Determination of IFN-α units

A stable reporter cell line using human retinal pigment epithelial cells (ATCC CRL2302) transfected with a plasmid containing interferon-stimulated response element promoter/enhancer elements driving a luciferase reporter gene. Cells were grown for 24 h before adding serial dilutions of recombinant IFN-α subtypes and a predetermined IFN-α lab standard for 6 h. The cells were lysed with Pierce Firefly Luc One-Step Assay Buffer (Thermofisher, Waltham, MA), and luciferase activity was measured 10 min later. The relative light units were plotted against the IFN-α dilutions using non-linear fit with three parameters and EC50 was determined. Units of IFN-α were calculated using EC50 values and known units/ μL of the standard.

### HIV-1 infection and IFN-α treatment in MT4C5 WT and KO cells

MT4C5 cells were kindly donated by Dr. Oliver Schwartz (Institut Pasteur, Paris, France). MT4C5 is a CD4^+^ T cell line expressing CCR5 and has little to no endogenous production of IFN-α(42). MT4C5 WT cells were maintained in RPMI-1640 (Hyclone, Logan, UT) supplemented with 10% FCS (Hyclone) and 1X PenStrep (Hyclone) (R-10) while KO made from MT4C5 cells were maintained with R-10 supplemented with 5μg/mL puromycin (Invivogen, San Diego, CA). Cells were either left untreated or treated with IFN-α2 or IFN-α14 (100U/mL, 500U/mL and 1000U/mL) for 24h and then infected with 0.1 MOI of HIV-1JR-CSF (credit proper people off AIDS Reagent) for 2 hours. Cells were then washed 2 times with complete media and plated in a 24 well plate. Post-infection all the cells were maintained in either IFN-α2 or IFN-α14 at the initial treatment dose.

### Guide RNA selection and cloning into lentiCRISPRv2

Each guide sequence was designed using Synthego’s CRISPR Design Tool. Guide sequences for MX2 (ENST00000330714), APOBEC3G (ENST00000407997) and ISG15 (ENST00000379389) were chosen from the list provided by Synthego’s CRISPR Design Tool (https://www.synthego.com/products/bioinformatics/crispr-design-tool). Location for cut site for MX2 (41.376.931, 41.377.150 and 41.377.009), APOBEC3G (39.080.934, 39.080.990 and 39.081.088) and ISG15 (1.014.247 and 1.014.006) were chosen such that the fragment that would be deleted would not be a multiple of three. DNA oligos (**Table 1**) for the guide sequences were ordered from Integrated DNA Technologies.

The protocol for cloning was adapted from Zhang Lab(51). The complementary oligos were phosphorylated using T4 PNK enzyme (New England Biolabs, Massachusetts, USA) and T4 Ligase buffer (Invitrogen, Carlsbad, California, USA) at 37°C for 1h. After phosphorylation, the complementary oligos were heated to 95°C in water bath and cooled down gradually to room temperature (22°C-25°C). LentiCRISPR v2 was a gift from Feng Zhang (Addgene plasmid # 52961 ; http://n2t.net/addgene:52961 ; RRID:Addgene_52961). LentiCRISPRv2 was digested using *BsmBI-v2* and the 12kb backbone was gel extracted using QIAquick Gel Extraction Kit Gel Purification (QIAGEN, Hilden, Germany). Oligos and the LentiCRISPRv2 backbone were ligated overnight at 16°C. Each of the ligated plasmids were transformed into *Stbl2* cells (Invitrogen) and were grown in freshly made LB Agar plates with 100μg/mL of ampicillin. Single colonies were grown in 5mL of LB media supplemented with 100μg/mL of ampicillin. Plasmids were isolated from transformed *Stbl2* cells using QIAprep Spin Miniprep Kit (QIAGEN) and PCR used to confirm the presence of the insert.

### Lentivirus production and transduction

HEK293 FT cells were maintained in DMEM (Gibco, Waltham, MA) supplemented with 10% FCS, 1X PenStrep, 1X Sodium Pyruvate and 1X NEAA (DMEM complete media). Cells were transfected with cloned LentiCRISPRv2 plasmid carrying oligos, pMD2.G and psPAX2 at a ratio of (1:2:1) using MEM (Gibco) and Lipofectamine 2000 (Invitrogen). After 72h of transfection, supernatant containing lentiviruses were collected and spun down at 1000 x g for 5mins to separate cell debris. Lentiviruses for same restriction factor were then incubated with MT4C5 cells for 24h. After 72h of transduction, freshly prepared selection medium (R-10 + 5 μg/mL puromycin) was added to the culture. After two weeks of selection, cells were treated with IFN-αs to confirm the KOs.

### Western blots

Cells were counted and 2X SDS buffer was added to cells. The mixture was then heated at 95°C for 10 mins and then spun at 13,000xg for 1 min. Samples were loaded onto a 12% SDS-PAGE gel for 1.2 hrs. After protein transfer was done onto the nitrocellulose membrane, blocking was done using 1X clear milk (Thermofisher). Membrane was then incubated with primary antibodies **(Table 2)** overnight at 4°C and washed 5 times with 0.1% tween-20 containing 1XPBS for 5mins each. Secondary antibodies **(Table 2)** were added and incubated at room temperature for one hour and then washed with 5 times with 0.1% tween-20 containing 1XPBS for 5mins each.

**Table 2.** Antibody dilutions.

| **<u>Sl. no.</u>** | **<u>Antibody</u>** | **<u>Host</u>** | **<u>Dilution</u>** | **<u>Primary/Secondary</u>** |
| --- | --- | --- | --- | --- |
| 1. | Hu-MX2 | Mouse | 1:100 | Primary |
| 2. | Hu-ISG15 | Mouse | 1:1000 | Primary |
| 3. | Hu-GAPDH | Rabbit | 1:10000 | Primary |
| 4. | m-IgG | Mouse | 1:10000 | Secondary |
| 5. | m-IgG | Rabbit | 1:10000 | Secondary |

### Genomic DNA isolation and PCR amplification

Genomic DNA from different experimental groups was extracted using the QIAamp DNA minikit (QIAGEN) kit, following the manufacturer’s instructions. The APOBEC3G sequence flanking the region targeted by guide RNAs was amplified using forward primer 5’- GGATGAAGCCTCACTTCAGAAA-3’ and reverse primer 5’- CACAGGTAAGTCTCATGCCGT-3’. Determination of total HIV-1 DNA was performed by quantitative real-time PCR using SsoAdvanced Universal SYBR Green Supermix (BIORAD, Hercules, CA) targeting the POL region of HIV-1 with forward primer 5’- ACTCCAGTATTTGCCATAAAGAAAAAA-3’ and reverse primer 5’- CTGTCTTTTTCTGGCAGCACTAT-3’ and GAPDH with forward primer 5’- GCTCTTAAAAAGTGCAGGGTCT-3’ and reverse primer 5’- GATGGTACATGACAAGGTGCG-3’.

### p24 enzyme linked immunosorbent assay

Cell supernatants were collected at day 6 for p24 ELISA (Advanced Bioscience Laboratories, Maryland, USA) following the manufacturer’s instructions using a laboratory optimized modification that halved the sample and reagent volumes used at each step. Plates were read in a 96 well plate reader at 450nm. Data acquisition was done using Softmax Pro (San Jose, CA).

### Infectivity Assay

TZM-bl reporter cells (NIH AIDS Research and Reference Reagent Program, from John Kappes, Xiaoyun Wu, and Tranzyme Inc.) were used to determine the number of infectious particles in the supernatant. Briefly, cells were plated 24h prior to the assay in a 96-well plate. Supernatants were 10X serially diluted and incubated with cells for 2h. Cells were then washed and cultured with fresh media for 48h. Cells were then fixed with 2% gluteraldehyde and X-gal solution was added and plate was incubated at 37°C at 5% CO2 for 24h. Spots were then counted at 4X magnification and the infectivity was normalised to the p24 of the supernatants.

### Detection of APOBEC3G signature mutations

Supernatants from MT4C5 WT or A3G KO treated with different IFN-α subtypes for 6 days were incubated with target cells (A3G KO) for 48h. The POL region was amplified, gel purified using QIAquick Gel Extraction Kit – Gel Purification (QIAGEN) and sent for sequencing (Plasmidsaurus, South San Francisco, CA) to determine the ratio of GG→ AG mutations vs total mutations.

### Statistics

Data were plotted with mean and standard deviation. Normality of each dataset was evaluated by Student’s unpaired t-test or one-way ANOVA with Tukey’s post-tests were performed accordingly and are specified within figure legends for each data set. P**<**0.05 was considered statistically significant. All statistics were performed using Prism 10 (GraphPad, San Diego, CA).

## ACKNOWLEDGEMENTS

Support for this work was provided by the Natural Sciences and Engineering Research Council of Canada (NSERC) and the Canadian Institutes of Health Research (CIHR).

